# The Small GTPase OsRac1 forms two distinct immune receptor complexes containing the PRR OsCERK1 and the NLR Pit

**DOI:** 10.1101/2020.07.01.183301

**Authors:** Akira Akamatsu, Masayuki Fujiwara, Satoshi Hamada, Megumi Wakabayashi, Ai Yao, Qiong Wang, Ken-ichi Kosami, Thu Thi Dang, Takako Kaneko-Kawano, Ko Shimamoto, Yoji Kawano

## Abstract

Plants employ two different types of immune receptors, cell surface pattern recognition receptors (PRRs) and intracellular nucleotide-binding and Leucine-rich repeat-containing proteins (NLRs), to cope with pathogen invasion. Both immune receptors often share similar downstream components and responses but it remains unknown whether a PRR and an NLR assemble into the same protein complex or two distinct receptor complexes. We have previously found that the small GTPase OsRac1 plays key roles in the signaling of OsCERK1, a PRR for fungal chitin, and of Pit, an NLR for rice blast fungus, and associates directly and indirectly with both of these immune receptors. In this study, using biochemical and bioimaging approaches, we reveal that OsRac1 formed two distinct receptor complexes with OsCERK1 and with Pit. Supporting this result, OsCERK1 and Pit utilized different transport systems for anchorage to the plasma membrane. Activation of OsCERK1 and Pit led to OsRac1 activation and, concomitantly, OsRac1 shifted from a small to a large protein complex fraction. We also found that the chaperone Hsp90 contributed to the proper transport of Pit to the plasma membrane and the immune induction of Pit. These findings illuminate how the PRR OsCERK1 and the NLR Pit orchestrate rice immunity through the small GTPase OsRac1.

## Introduction

Plants utilize two layers of immune response to cope with pathogen infection. The first layer is known as pathogen-/microbe-associated molecular pattern (PAMP/MAMP)-triggered immunity (PTI/MTI), while the second layer is called effector-triggered immunity (ETI) (Dangl et al. 2013; Dodds and Rathjen 2010). PTI is triggered by transmembrane pattern recognition receptors (PRRs) and induces early responses (Couto and Zipfel 2016). Most PRRs are categorized into two protein families consisting of receptor-like kinases (RLKs) and receptor-like proteins (RLPs) (Monaghan and Zipfel 2012). RLKs perceive signals through their extracellular domains, transmit these signals to kinase domains, and phosphorylate their downstream intracellular signaling molecules. ETI is initiated by either direct or indirect recognition of pathogen effectors by the nucleotide-binding domain and leucine-rich repeats (NLR) family proteins (Cui et al. 2015).

In at least some cases, PTI and ETI employ similar signaling machinery such as Ca^2+^ signaling, reactive oxygen species (ROS) generation, transcriptional reprogramming, and MAP kinase (MAPK) cascade activation (Peng et al. 2018; Tsuda et al. 2013). Transcriptome analysis has revealed that the sets of genes induced by PTI and ETI overlap. However, the immune responses induced by ETI are generally more rapid, prolonged, and robust than those induced by PTI (Dodds and Rathjen 2010; Tao et al. 2003; Thomma et al. 2011; Tsuda and Katagiri 2010). Moreover, PRRs and NLRs require each other to effect robust disease resistance (N’gou et al. 2020; Yuan et al. 2020). These results suggest that PTI and ETI share the same or similar signaling machinery, while their dynamics and strength are different. Qi et al. previously demonstrated that an *Arabidopsis* PRR, FLAGELLIN-SENSING 2 (FLS2), is physically associated with three plasma membrane (PM)-localized NLR proteins, RPS2, RPM1, and RPS5 (Qi et al. 2011). However, it is currently unclear whether physical interaction between PRRs and NLRs is a general feature and how two different types of immune receptors, PRRs and NLRs, induce similar responses.

Heat shock proteins (Hsps) are abundant and highly conserved proteins that accumulate in response to various stresses and serve as molecular chaperones for diverse client proteins. Hsp90 associates with many co-chaperones and cofactors to promote proper folding and maturation of client proteins (Pearl and Prodromou 2006). Previous studies have shown that Hsp90 plays critical roles in NLR functions. Indeed, suppression of HSP90 function leads to increased susceptibility to pathogens (Hubert et al. 2003; Lu et al. 2003). Hsp90 forms a complex(es) with co-chaperones such as Suppressor of G2 allele of skp1 (SGT1) and required for Mla12 resistance1 (RAR1) (Shirasu et al. 1999; Takahashi et al. 2003), and contributes to the stabilization of NLR proteins (Kadota and Shirasu 2012). Hsp90.7, an ER-localized Hsp, is required for the correct folding and/or complex formation of the two RLKs CLAVATA 1 and 2 to control shoot and floral meristem development (Ishiguro et al. 2002).

Members of the small GTPase Rac/Rop family act as molecular switches and play crucial roles in a variety of plant physiological processes (Berken 2006; Nibau et al. 2006). The small GTPase OsRac1 functions as a key regulator in both PTI and ETI in rice (Kawano et al. 2010b; Kawano et al. 2014b; Kawano and Shimamoto 2013). OsRac1 contributes to PTI triggered by two elicitors, chitin and sphingolipid, derived from fungal pathogens. We have also revealed that an OsCERK1–OsRacGEF1–OsRac1 module is involved in early signaling for chitin-induced immunity (Akamatsu et al. 2015; Akamatsu et al. 2013). After sensing chitin, the chitin receptor complex containing the RLP OsCEBiP and the RLK OsCERK1 phosphorylates OsRacGEF1, which is a PRONE family activator protein of OsRac1. OsCERK1-dependent phosphorylation of OsRacGEF1 leads to OsRac1 activation, resulting in the induction of immune responses. Hsp90 and its co-chaperone Hop/Sti1 complex contribute to the maturation and intracellular transport of the OsCERK1 complex (Chen et al. 2010a). Moreover, OsRac1 also forms a complex(es) with various proteins including Hsp70, the scaffold protein OsRACK1, the lignin biosynthesis enzyme OsCCR1, and OsMPK6 (Kawasaki et al. 2006; Kim et al. 2012; Lieberherr et al. 2005; Nakashima et al. 2008; Thao et al. 2007). In ETI, NLR proteins employ a different mechanism to elicit OsRac1 activation from PRRs (Kawano et al. 2010a; Wang et al. 2018). Two NLR proteins, Pit and Pia, for rice blast fungus directly bind to OsSPK1, which is a DOCK family activator protein for OsRac1, and induce OsRac1 activation through OsSPK1, leading to disease resistance to rice blast fungus (Kawano et al. 2014a; Ono et al. 2001; Wang et al. 2018). So far, many players get involved in the immune complex(es) with OsRac1, however which components form a distinct complex(es) in PTI and ETI as well as their functions remain to be explored.

In this study, we demonstrated that OsRac1 formed two distinct immune receptor complexes with the RLK OsCERK1 and the NLR Pit. Chitin perception or induction of an active form of Pit made OsRac1 into an active form, which led to the redistribution of OsRac1 from a low to a high molecular weight complex. Hsp90 appears to play a critical role in the proper localization of Pit. These results shed light on the underlying molecular mechanisms of how PRRs and NLRs orchestrate rice immunity through the small GTPase OsRac1.

## Results

### Two distinct immune receptor complexes: the PRR OsCERK1 and the NLR Pit

The small GTPase OsRac1 functions as a downstream molecular switch for two different types of immune receptors, the PRR OsCERK1 and the NLR Pit (Akamatsu et al. 2013; Kawano et al. 2010a; Wang et al. 2018), and we therefore wondered whether the three proteins form a ternary complex or two distinct complexes. We performed an immunoprecipitation assay using rice suspension cells expressing *Myc-OsRac1* with *OsCERK1-FLAG* and/or *Pit-HA*. When OsCERK1-FLAG or Pit-HA was immunoprecipitated, OsRac1 coprecipitated with each (Fig. 1A). However, we observed no interaction between OsCERK1 and Pit even when we reciprocally precipitated both receptors, implying that OsCERK1_and Pit form two distinct immune receptor complexes with OsRac1. To validate this result in living cells, we employed two bioimaging methods. First, we tested for an interaction between OsCERK1 and Pit *in vivo* using bimolecular fluorescence complementation (BiFC) assays. To quantify the interactions in BiFC assays, we measured the frequency of reconstituted Venus-positive protoplasts in each combination of constructs. When OsCERK1 tagged with the N-terminal domain (aa 1-154) of Venus (OsCERK1-Vn) and Pit tagged with the C-terminal domain (aa 155-238) of Venus (Pit-Vc) were co-expressed in rice protoplasts, Venus fluorescence was not detected under conditions in which the known interactions between OsRac1 and Pit as well as OsCERK1 and Hop/Sti1 were confirmed (Chen et al. 2010a; Kawano et al. 2010a) (Fig. 1B). Since we have previously shown that both Pit and OsRac1 are localized at the plasma membrane through palmitoylation, a lipid modification (Chen et al. 2010b; Kawano et al. 2014a; Ono et al. 2001), we further tested the distribution of OsCERK1 and Pit in living cells using variable angle epifluorescence microscopy, also called variable incidence angle fluorescence microscopy (VIAFM) (Fujimoto et al. 2010; Konopka and Bednarek 2008). VIAFM is a derivative of total internal reflection fluorescence microscopy employing an evanescent wave that excites fluorescent proteins selectively in a region of the specimen beneath the glass-water interface, such as the plasma membrane and the cytoplasmic zone immediately beneath the plasma membrane of cells. We transfected rice protoplasts with *OsCERK1-mCherry* and *Pit-mGFP* vectors and observed the localization of the expressed proteins (Fig. 1C). OsCERK1-mCherry and Pit-mGFP showed small fluorescent particles, whereas there were no detectable particles in control mGFP-expressing cells (Supplemental Fig. 1D). The result of dual-fluorescence imaging using OsCERK1-mCherry and Pit-WT-mGFP clearly demonstrated that almost none of the OsCERK1-mCherry and Pit-mCherry particles overlapped with each other (Fig. 1C). On the other hand, OsCERK1-mCherry co-localized well with mEGFP-OsCEBiP, a co-chitin receptor (Supplemental Fig. 1E–G). Taken together, these results indicate that OsCERK1 and Pit form two different immune complexes with OsRac1.

**Fig. 1.**
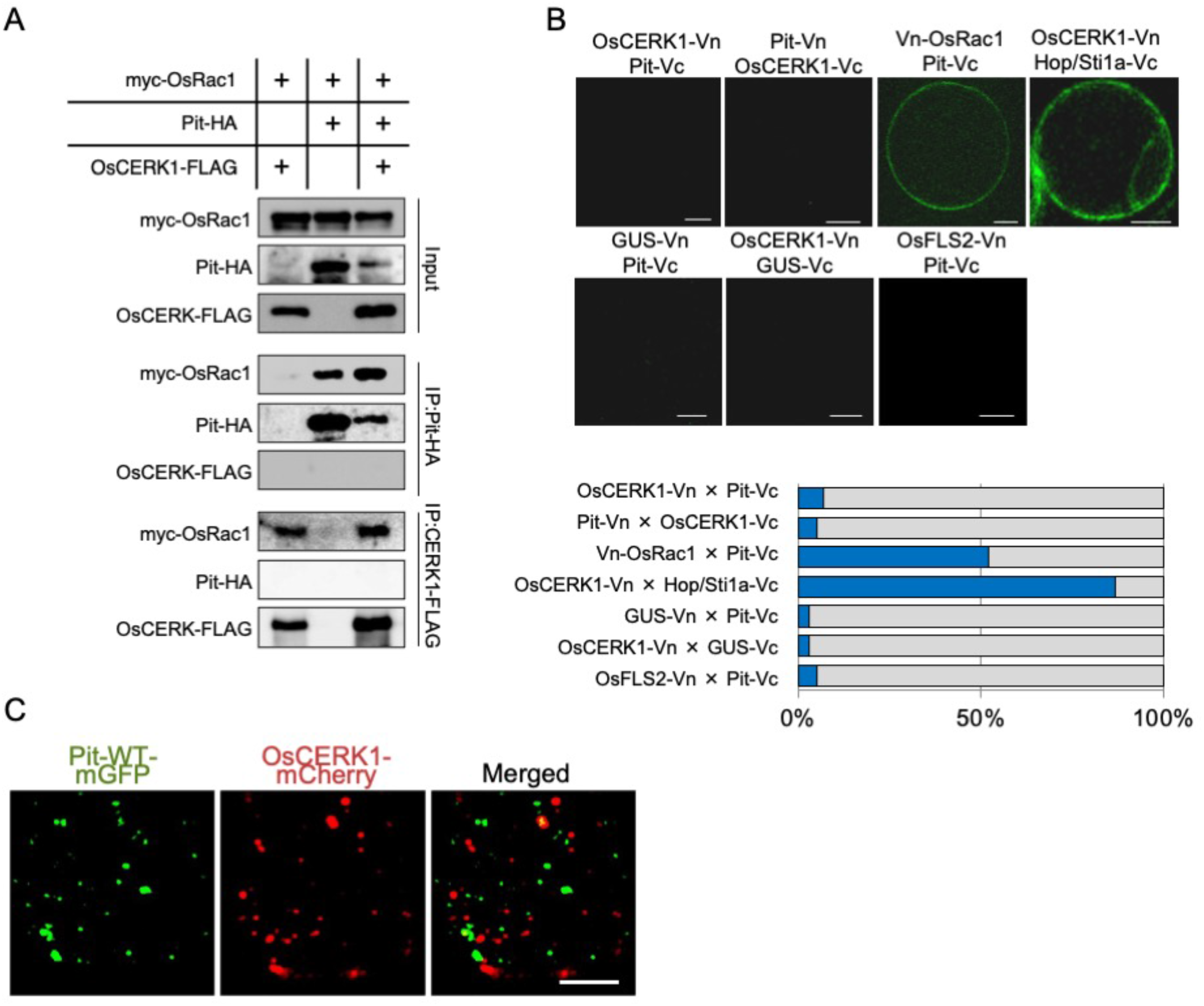
OsRac1 forms two distinct immune receptor complexes. (A) *In vivo* interaction between the chitin receptor OsCERK1 and the NLR protein Pit. Co-IP was performed using an anti-HA and anti-FLAG antibodies, and the proteins were detected by immunoblot with the indicated antibodies. (B) BiFC assay between OsCERK1 and Pit. Expression of the indicated genes was driven by the CaMV 35S promoter. The graph shows the percentage of BiFC positive cells. Scale bars, 5 μm. (C) Representative VIAFM images of rice protoplasts expressing Pit-WT-mEGFP and OsCERK1-mCherry. Left, center, and right panels are GFP, mCherry, and merged images, respectively. Scale bar, 5 μm.

### Different signaling components and intracellular transport of OsCERK1 and Pit

Based on a number of protein–protein interactions and functional studies, we have previously demonstrated that OsCERK1 directly binds to chaperone Hsp90 and co-chaperone Hop/Sti1a (Chen et al. 2010a). Moreover, OsRac1 interacts either directly or indirectly with Hsp90, its co-chaperones Hop/Sti1a, SGT1, and RAR1, the scaffold protein RACK1A, and MAP kinase MPK6, and these components play important roles in both PTI and ETI (Akamatsu et al. 2013; Chen et al. 2010a; Kawano et al. 2010a; Lieberherr et al. 2005; Nakashima et al. 2008; Thao et al. 2007). To identify the components of OsCERK1 and Pit complexes, we performed an immunoprecipitation assay using rice suspension cells. The two different immune receptor complexes contain shared components including Hsp90, Hop/Sti1, and OsRac1, but they do not include the OsRac1 interactors RACK1A, RAR1, or MPK6 (Fig. 2A and 2B). Interestingly, SGT1 is a specific component in the Pit complex (Fig. 2B).

**Fig. 2.**
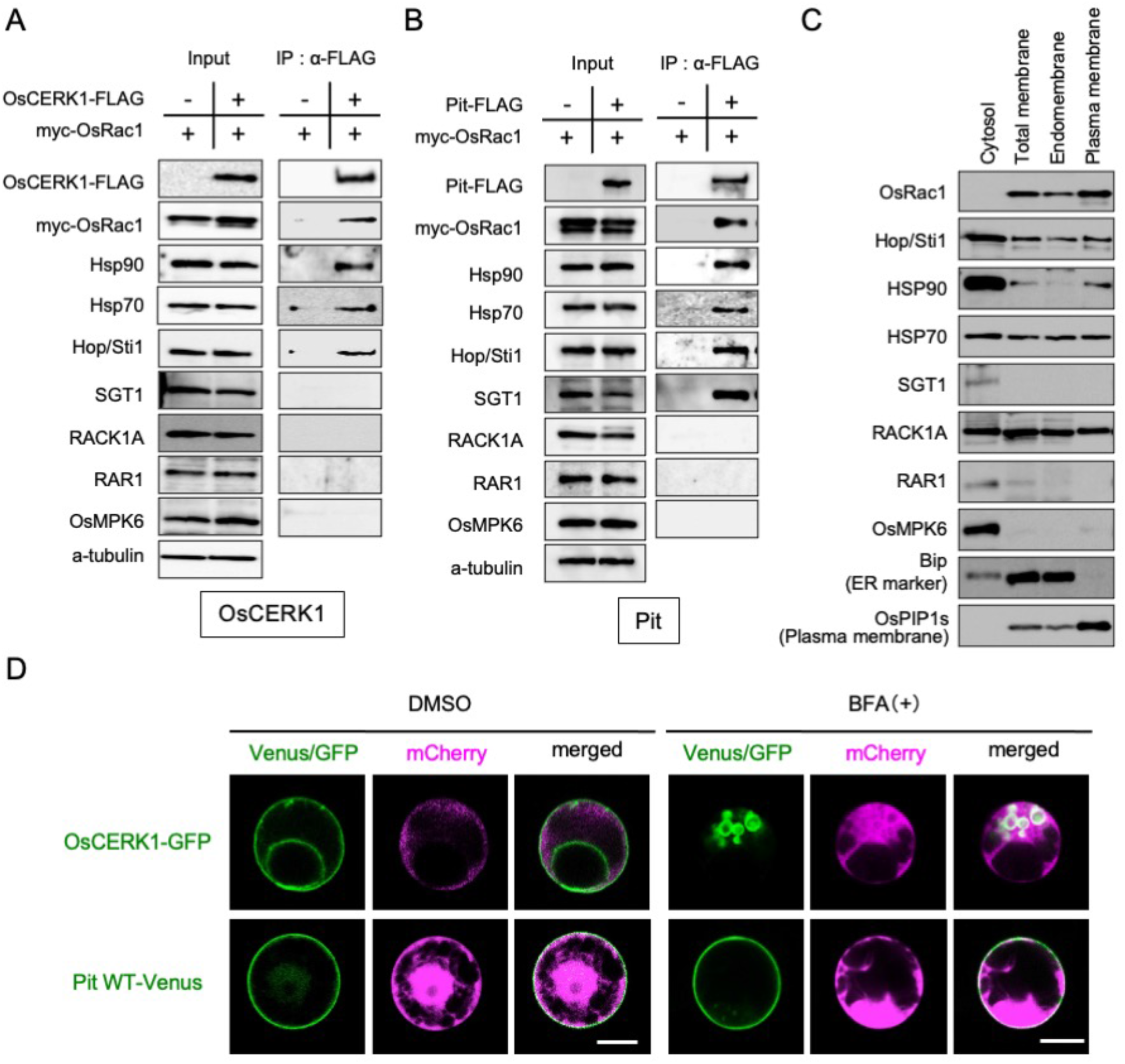
Components of OsCERK1- and Pit-containing immune complexes. (A, B) Co-IP of (A) OsCERK1- and (B) Pit-containing immune complexes. Co-IP was performed using anti-FLAG antibody, and the proteins were detected by immunoblot with the indicated antibodies. (C) Distribution of defense-related proteins. An aqueous two-phase partitioning experiment was performed and the proteins were detected by immunoblot with the indicated antibodies. (D) Localization of OsCERK1 and Pit in rice protoplasts in the presence of BFA. OsCERK1-GFP or Pit1-Venus was co-transfected with mCherry. Sixteen hours after BFA treatment, the transfected cells were observed under a microscope. Scale bars, 5 μm.

Generally, small GTPase Rac/Rop family proteins are localized at the plasma membrane as a result of post-translational modification (Ono et al. 2001; Yalovsky et al. 2008). We have previously shown that OsRac1 localizes predominantly at the PM (Chen et al. 2010a), and that Hop/Sti1a and Hsp90 are present in the PM-rich fraction (Chen et al. 2010b). To more precisely examine the intracellular distribution of the components, we performed an aqueous two-phase partitioning experiment and found that they show three different patterns of distribution. OsRac1 localized in the plasma membrane and endomembrane fractions. Hop/Sti1a, Hsp70, and RACK1A were dispersed in the cytosol, endomembrane, and plasma membrane fractions. In contrast, Hsp90, SGT1, RAR1, and OsMPK6 were restricted mainly to the cytosol fraction, although a small proportion of Hsp90 partitioned to the plasma membrane fraction (Fig. 2C).

Next, we compared the intracellular transport system of OsCERK1 with that of Pit in rice protoplasts. We have previously revealed that OsCERK1 is sensitive to brefeldin A (BFA), an inhibitor of anterograde endoplasmic reticulum–Golgi transport, and is transported by a small GTPase Sar1-dependent vesicle trafficking pathway (Chen et al. 2010a; Takeuchi et al. 2000) (Fig. 2D). In contrast, Pit is a palmitoylated protein and localizes at the plasma membrane in rice protoplasts (Kawano et al. 2014a), and here we revealed that Pit is insensitive to BFA (Fig. 2D), indicating that OsCERK1 and Pit employ different intracellular transport pathways to reach the plasma membrane.

### Monitoring the OsRac1 complex during OsCERK1-triggered immunity

To monitor the time course of OsRac1 activation after treatment with the fungal PAMP chitin, we employed a GST-PAK CRIB pull-down assay. This method exploits the Cdc42/Rac interactive binding (CRIB) domain of the Rac effector PAK1 (PAK CRIB), which shows a high affinity only for the active GTP-bound form of Rac, and not for the inactive GDP-bound form. This feature provides a useful tool to monitor the activation state of OsRac1 *in vivo* (Kawano et al. 2010a; Sander et al. 1998). As shown in Fig. 3A, a constitutively active mutant of OsRac1 (CA-OsRac1) specifically bound to PAK CRIB but a dominant-negative mutant (DN-OsRac1) did not, indicating that PAK CRIB should efficiently isolate the active GTP-bound form of OsRac1 from the crude cell lysate of suspension cells. Next, therefore, we prepared rice suspension cells expressing *myc-OsRac1 WT* to monitor the OsRac1 activation state after chitin treatment. A pull-down assay revealed statistically significant OsRac1 activation, beginning by 10 min after chitin treatment and lasting until at least 60 min (Fig. 3B). We investigated the dynamics of the OsRac1 complex by gel filtration and found that OsRac1 was divided into two groups: the high molecular weight OsRac1 fractions (HOR) (fractions 23–25; about 300 kDa) and the low molecular weight OsRac1 fractions (LOR) (fractions 29–31; about 50 kDa) (Fig. 3C). Intriguingly, a shift of WT OsRac1 from LOR to HOR was observed after a 10-min chitin treatment. We compared activation levels of OsRac1 between LOR and HOR using a GST-PAK CRIB pull-down assay. In the absence of chitin, total OsRac1 was distributed predominantly in LOR and gradually moved to HOR after chitin treatment (Fig. 3D, lower panel). Concomitantly, the active GTP form of OsRac1 was increased in HOR_(Fig. 3D, higher panel), implying that activation of OsRac1 promotes the shift to HOR. To test this hypothesis, we carried out a gel filtration assay using rice suspension cells expressing CA-OsRac1 and DN-OsRac1 (Fig. 3E). CA-OsRac1 was distributed exclusively in HOR, while DN-OsRac1 existed mainly in LOR. Next, we measured the amount of OsRac1 in the OsCERK1 complex after chitin treatment and found that chitin treatment induced the dissociation of OsRac1 from OsCERK1, but there were no obvious changes in the other components (Fig. 3F).

**Fig. 3.**
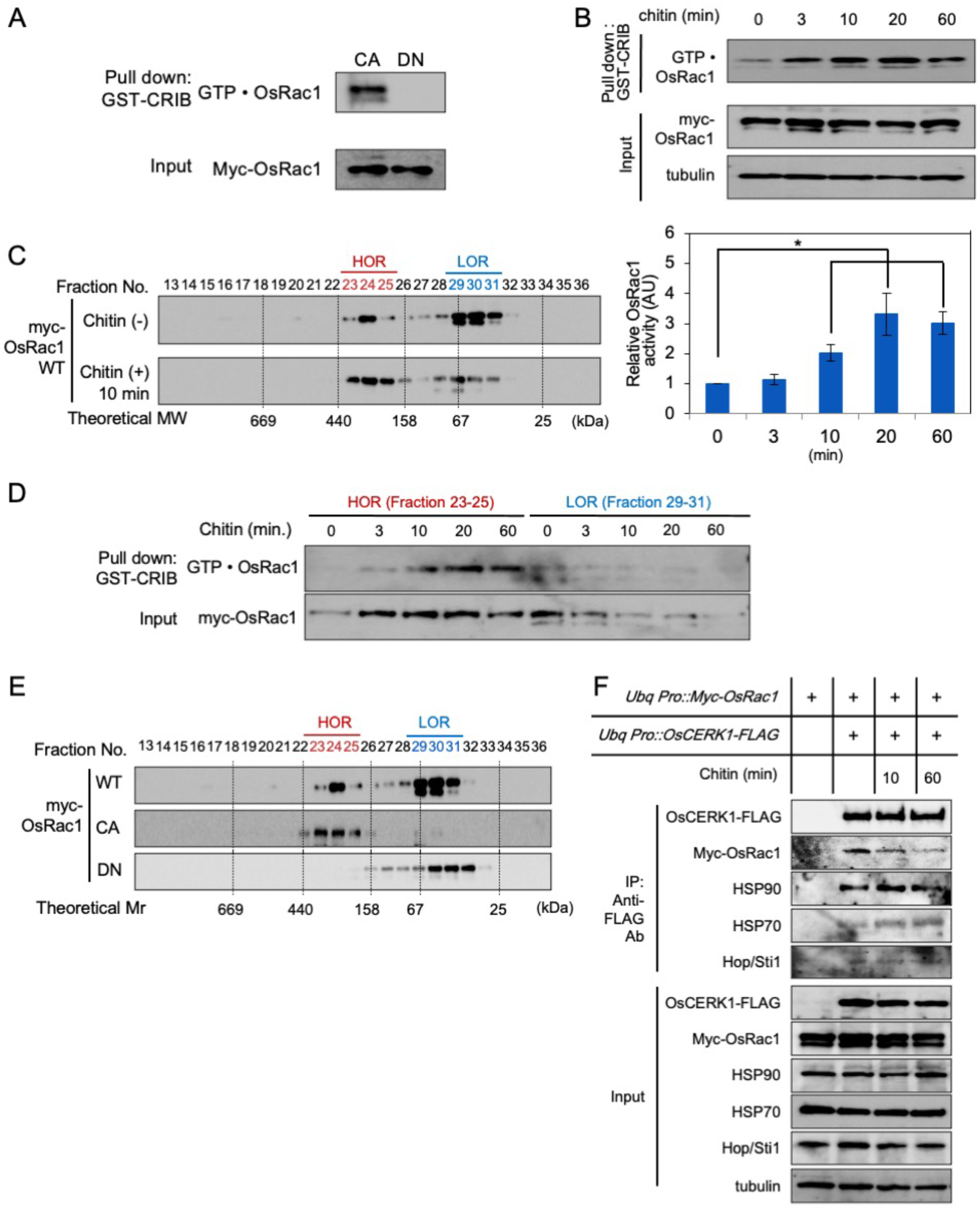
Active OsRac1 forms a large immune complex after chitin treatment. (A) GST-PAK CRIB pull-down assay using rice suspension cells expressing a dominant-negative mutant of *OsRac1* (*DN-OsRac1*) and a constitutively active mutant of *OsRac1* (*CA-OsRac1*). The band of GTPoOsRac1 indicates the amount of the active form of OsRac1. (B) Monitoring OsRac1 activation after chitin treatment. Rice suspension cells expressing myc-OsRac1 WT were treated with chitin and the resultant cell lysates were subjected to GST-PAK CRIB pull-down assay to detect OsRac1 activation. The graph indicates the band intensity analyzed by ImageJ software. Error bars indicate the SD. Different letters above bars indicate a significant difference determined by Student’s *ř*-test (*P* < 0.05). (C) Gel filtration fractions of protein extracts from rice suspension cells expressing *OsRac1 WT* before and after chitin treatment (upper and lower panels) were subjected to immunoblot analyses using an anti-Myc antibody. Fraction numbers and relative molecular masses (kDa) are indicated at the top and bottom, respectively. (D) Combined high-molecular-weight OsRac1 fractions (HOR) (fractions 23–25 in (C)) or low-molecular-weight OsRac1 fractions (LOR) (fractions 29– 31) were applied to GST-PAK CRIB pull-down assay to monitor OsRac1 activation. (E) Gel filtration fractions of protein extracts from rice suspension cells expressing *OsRac1 WT, CA, and DN*. Fraction numbers and relative_molecular masses (kD) are indicated at the top and bottom, respectively. (F) Components of the OsCERK1 complex after chitin treatment. OsCERK1-FLAG was immunoprecipitated with an anti-FLAG antibody. The precipitates were immunoblotted with the indicated antibodies.

### Monitoring the OsRac1 complex during Pit-triggered immunity

The active form of Pit activates OsRac1, and this activation seems to be critical for the induction of disease resistance to rice blast fungus (Kawano et al. 2010a; Wang et al. 2018). To examine the dynamics of the Pit complex, we generated rice suspension cells expressing *myc-OsRac1 WT* and either *Pit WT-FLAG* or *Pit D485V-FLAG*, which is a constitutively active mutant and triggers OsRac1 activation and cell death without fungus infection, under control of an estradiol-inducible promoter. We first checked the induction of *Pit D485V-FLAG* by estradiol at the RNA and protein levels (Fig. 4A). *Pit D485V-FLAG* mRNA was detected by RT-PCR after 1 h of estradiol treatment and gradually increased until 16 h. Correspondingly, a very faint band of Pit D485V-FLAG protein was observed 4 h after estradiol treatment began, and th protein had accumulated by 8 h. To analyze OsRac1 activation by Pit, we performed GST-pull down using GST-PAK CRIB (Fig. 4B). Induction of Pit D485V-FLAG by estradiol triggered activation of OsRac1; in contrast, induction of Pit WT-FLAG did not activate OsRac1 (Fig. 4B). Consistent with this, the transcript level of the defense gene *PAL1* in the Pit D485V cell line increased upon estradiol treatment (Fig. 4C). Similar to OsRac1 dynamics after chitin treatment, the ratio of HOR to LOR after the addition of estradiol was higher than that before the estradiol treatment, suggesting that OsRac1 shifts from LOR to HOR as a consequence of the expression of *Pit D485V* (Fig. 4D). Next, we tested whether chitin treatment affects OsRac1–Pit interaction and found that OsRac1 was dissociated from Pit by the addition of chitin (Fig. 4E), implying that there is crosstalk between OsCERK1 signaling and Pit signaling.

**Fig. 4.**
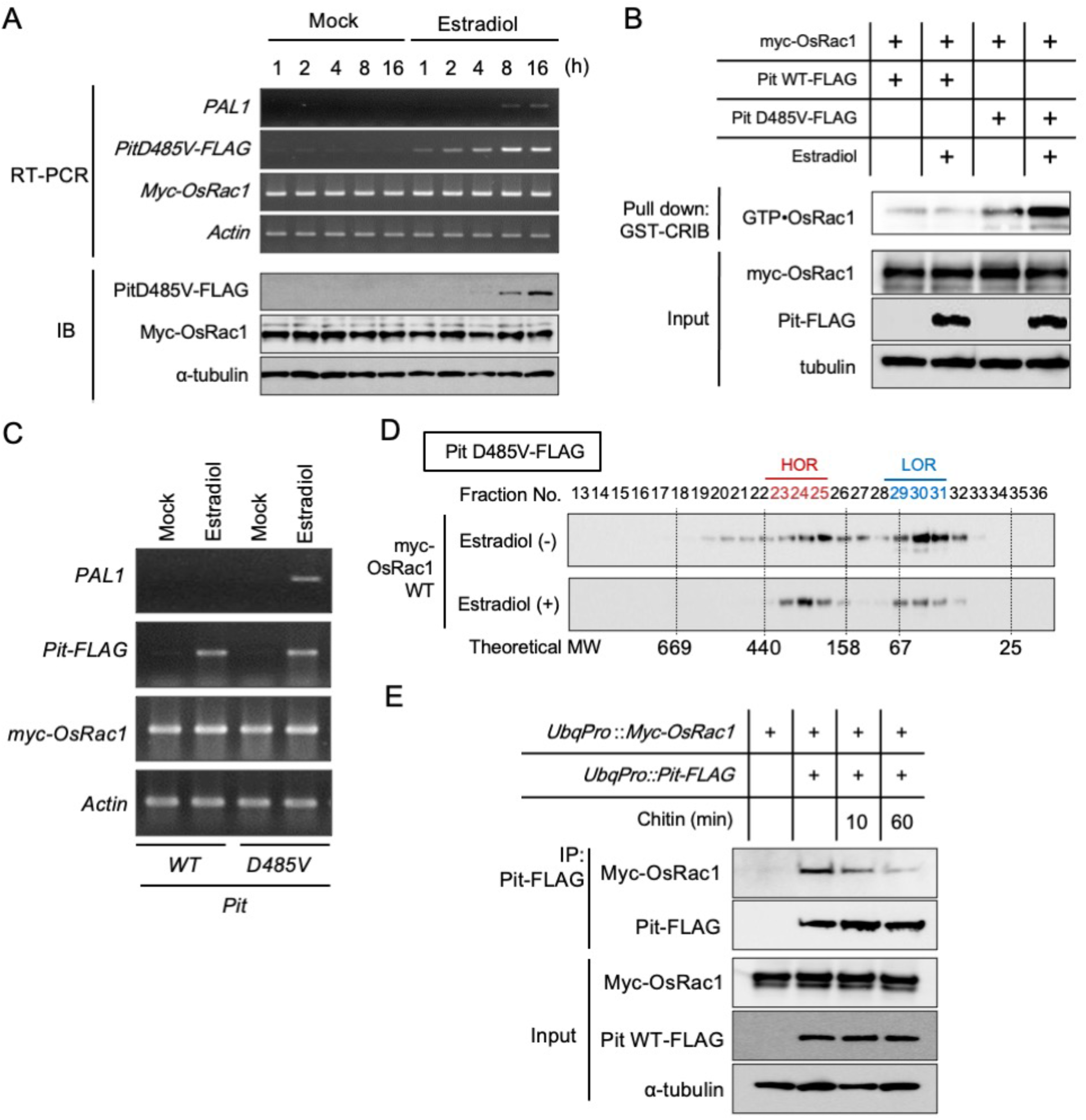
Active-form Pit shifts OsRac1 to the larger immune complex. (A) Induction of constitutively active *Pit* (*Pit D485V*) mRNA and protein by estradiol treatment. (B) Expression of Pit D485V triggers OsRac1 activation. After the induction of Pit D485V by estradiol, we carried out a GST-PAK CRIB pull-down assay to detect OsRac1 activation. (C) Defense gene *PAL1* is induced by the expression of *Pit D485V*. (D) Gel filtration fractions of protein extract from rice suspension cells expressing OsRac1 WT before and after Pit D485V induction. Fraction numbers and relative molecular masses (kD) are indicated at the top and bottom, respectively. (E) Interaction between Pit and OsRac1 after chitin treatment. After chitin treatment, Pit-FLAG was precipitated by an anti-FLAG antibody. The resultant precipitates were immunoblotted with anti-Myc antibody.

### Hsp90 is an essential component of Pit-dependent immunity

Since Hsp90 is critical for stabilizing several NLR proteins (Hubert et al. 2003; Kadota and Shirasu 2012; Takahashi et al. 2003), its roles in ETI have been determined using an Hsp90-specific inhibitor, geldanamycin (GDA). To clarify the role of Hsp90 in Pit-induced defense responses, we tested the effect of GDA on Pit D485V-induced cell death and ROS production in *Nicotiana benthamiana*. Overexpression of the constitutively active mutant Pit D485V triggered cell death and ROS production, but this effect was suppressed by the co-infiltration of GDA (Fig. 5A) and the knockdown of endogenous *NbHsp90* by virus-induced gene silencing (Fig. 5B). Consistent with these observations, *PAL1* induction by Pit D485V-FLAG was attenuated by GDA treatment in rice suspension cells (Fig. 5C), indicating that proper Hsp90 activity is required for Pit-induced immunity.

**Fig. 5.**
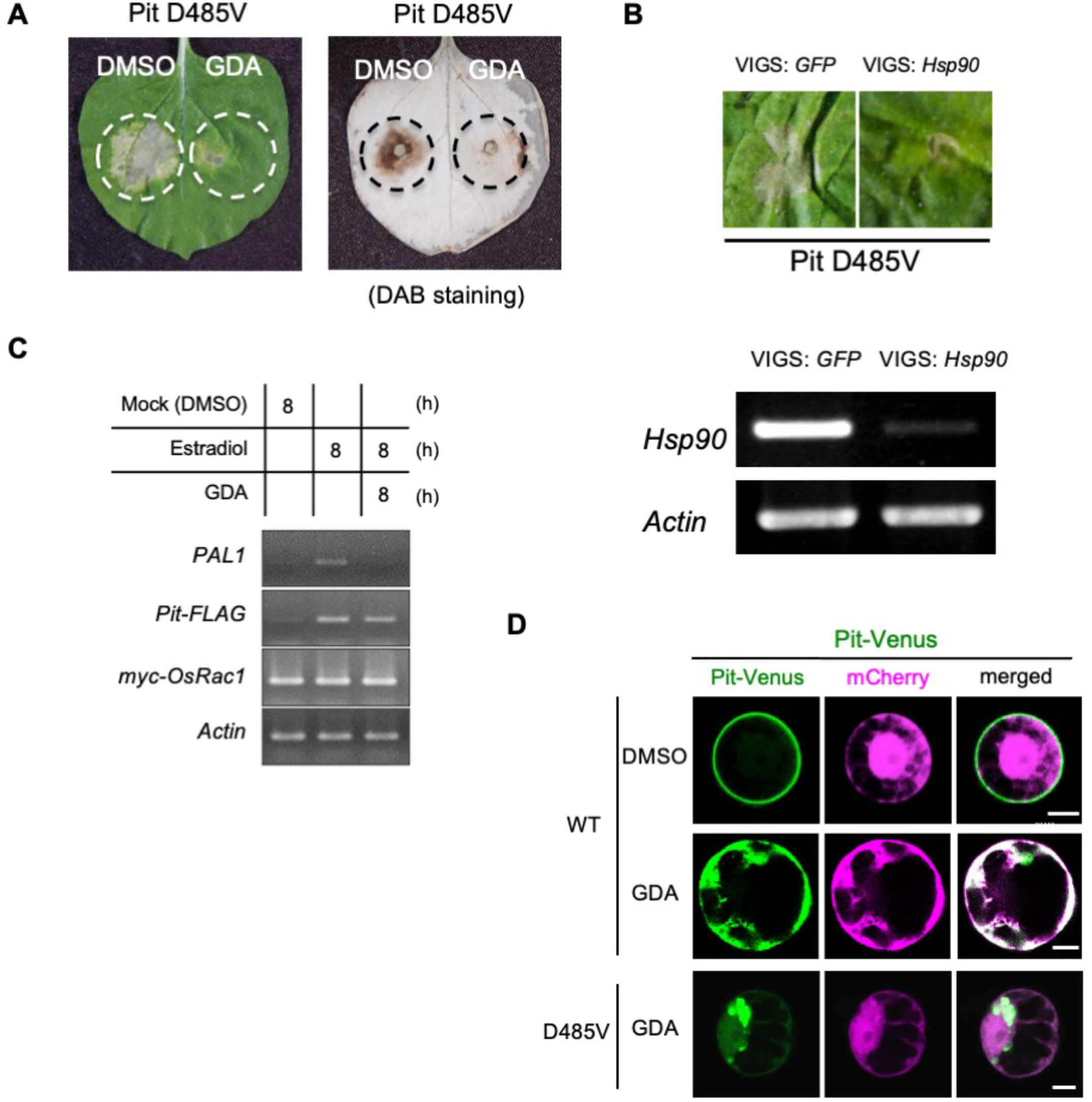
Hsp90 contributes to Pit-induced immunity. (A) Suppression of Pit-triggered cell death and ROS production by the Hsp90 inhibitor GDA. In the presence or absence of GDA, Pit D485V-induced cell death (left image) and ROS (right image) were examined in *N. benthamiana* leaves. (B) Suppression of Pit D485V-induced cell death by virus-induced gene silencing (VIGS) of Hsp90. *N. benthamiana* plants were inoculated with pGPVX:GFP or pGPVX:Hsp90 (10-186), and three weeks later the upper leaves were infiltrated with a mixture of *Agrobacterium* cultures carrying pGWB2-Pit D485V transgenes. Cell death developed by 7 days after inoculation (upper panels). mRNA expression of *Hsp90* and the internal control *Actin* was detected by RT-PCR (lower panels). (C) Inhibition of *Pit D485V-* induced *PAL1* expression by treatment of with GDA. mRNA expression of *PAL1*, *Pit*, *OsRac1*, *Hsp90*, and *Actin* was detected by RT-PCR. (D) Localization of Pit-Venus in rice protoplasts in the presence of GDA. Pit-Venus was co-transfected with mCherry. Twelve hours after GDA treatment, the transfected cells were observed under a microscope. Scale bars, 5 μm.

Finally, we monitored the localization of Pit in the presence of GDA in rice protoplasts. As we have reported previously, Pit WT-Venus was localized in the plasma membrane, but the addition of GDA abolished this plasma membrane localization of Pit WT and it accumulated instead in the cytosol (Fig. 5D). We could not observe a fluorescent signal of the constitutively active mutant Pit D485V-Venus in the absence of GDA, probably due to cell death. Interestingly, a clear fluorescent signal of Pit D485V-Venus was detected at the perinuclear region in the presence of GDA. This result implies that Hsp90 contributes to he maturation and/or proper plasma membrane localization of Pit and that it is indispensable for Pit’s function.

## Discussion

### OsRac1 is a component of two distinct receptor complexes

OsRac1 is one of the critical regulators in rice immunity, working with two different types of immune receptors, the PRR OsCERK1 and the NLR Pit (Akamatsu et al. 2013; Kawano et al. 2010a; Wang et al. 2018). Consistent with this, we here showed that OsRac1 was associated with both OsCERK1 and Pit but formed two distinct receptor complexes (Fig. 1 and 2). It appears that OsRac1 does not interact directly with OsCERK1 but requires a mediator protein, Hop/Sti1, to bind to OsCERK1 (Chen et al. 2010a). In contrast, OsRac1 associates directly with the NB-ARC domain of Pit (Kawano et al. 2010a). In general, PTI and ETI employ common signaling pathways such as ROS and the MAPK cascade, but immune responses by ETI are more robust and prolonged than those by PTI (Tsuda and Katagiri 2010). OsRac1 may be one of the common key machineries that control both PTI and ETI in rice. OsRac1 regulates ROS production in PTI and ETI, possibly through direct interaction with the NADPH oxidases RbohB/H (Kawasaki et al. 1999; Kosami et al. 2014; Nagano et al. 2016; Wong et al. 2007). OsRac1 forms a complex with and controls OsMPK6 at the protein level (Lieberherr et al. 2005). Further studies are needed to elucidate how OsRac1 contributes to PTI and ETI in a mechanistically different manner.

In this study, we revealed that although both OsCERK1 and Pit were localized in the plasma membrane (Fig. 2D), they utilize different transport systems to anchor themselves to the plasma membrane. We previously found that cysteine 97 and 98 in the N-terminal CC region of Pit are palmitoylation sites that play critical roles in its membrane localization and interaction with OsRac1 (Kawano et al. 2014a). Palmitoylation, also known as S-acylation, is the reversible post-translational addition of fatty acids to proteins and serves to target proteins to specific membrane compartments and/or microdomains (Hemsley 2015). OsCERK1 depends on COPII-mediated ER-to-Golgi traffic and on the *trans-*Golgi network for its transport to the plasma membrane (Fig. 2D) (Akamatsu et al. 2013; Chen et al. 2010a). Our previous BiFC analyses imply that OsRac1 is associated with OsCERK1 and Pit in different places: OsRac1 forms a complex with OsCERK1 through Hop/Sti1 possibly in the ER (Chen et al. 2010a) and with Pit at the plasma membrane (Kawano et al. 2014a), supporting our new observation that OsRac1 participates in distinct OsCERK1- and Pit-containing immune receptor complexes (Fig. 2).

### OsRac1 assembles into large protein complexes during PTI and ETI

Gel filtration and pull-down assay using GST-PAK CRIB revealed that the activation of OsCERK1 and Pit led in turn to OsRac1 activation, which induced a shift of OsRac1 from the LOR to the HOR, suggesting that OsRac1 activation by OsCERK1 and Pit activation assembles in the large protein complexes. OsRac1 belongs to the Rac/Rop family of small GTPases, which function as a molecular switch by cycling between GDP-bound inactive and GTP-bound active forms in cells (Kawano et al. 2014b). The active GTP-bound form of Rac/Rop binds to downstream target proteins to control various cellular events (Kawano et al. 2014b). Until now, we have identified various direct downstream target proteins of OsRac1, including the NADPH oxidases RbohB/H (Kosami et al. 2014; Nagano et al. 2016; Wong et al. 2007), the lignin biosynthesis key enzyme cinnamoyl-CoA reductase (Kawasaki et al. 2006), the co-chaperones Hop/Sti1, and the scaffold protein RACK1 (Nakashima et al. 2008). The direct binding of OsRac1 to these downstream target proteins probably causes the shift of OsRac1 from the LOR to the HOR. We previously proposed that the OsCERK1–OsRacGEF1–OsRac1 module is one of the key components in chitin signaling in rice (Akamatsu et al. 2013). Here, we observed that the majority of OsRac1 existed in the LOR in the absence of chitin, and we found no obvious increment of OsRac1 protein in the OsCERK1 complex after chitin treatment, implying that the association of OsRac1 with OsRacGEF1 in the OsCERK1 complex is transient.

### Roles of chaperones and co-chaperones in plant immunity

Here, we revealed that both OsCERK1 and Pit are also associated with the core chaperones Hsp90 and Hsp70 and the co-chaperone Hop/Sti1, and that SGT1 is a specific component of the Pit complex (Fig. 2A and 2B). We previously found that Hsp90 and Hop/Sti1 directly bind to OsCERK1 at the endoplasmic reticulum and contribute to the maturation of OsCERK1 and its transport to the plasma membrane (Chen et al. 2010a). Interactions between Hsp90 and various NLR proteins including RPM1, N, MLA1, MLA6, and Bs2 have been reported, and the LRR domain is likely an important site for NLR protein binding to Hsp90’? (Bieri et al. 2004; Hubert et al. 2003; Leister et al. 2005; Liu et al. 2004). One of the major roles for the Hsp90–SGT1–RAR1 complex is apparently to stabilize NLR proteins. GDA treatment or knockdown or knockout of *Hsp90, SGT1*, and *RAR1* compromises the plant’s disease resistance to pathogens and reduces the levels of NLR proteins (Kadota et al. 2010). The complex presumably controls the active/inactive state of NLR proteins. The pepper NLR protein Bs2 an intramolecular interaction between NB and LRR domains and this intramolecular interaction_that was abolished’ by silencing *SGT1* (Leister et al. 2005), implying that SGT1 participates in the intramolecular interactions within NLR proteins. In this study, we revealed that the attenuation of Hsp90 expression or function compromised Pit-induced immune responses. Moreover, GDA treatment perturbed the plasma membrane localization of Pit (Fig. 5D). Taken together, these results suggest that the proper function of Pit requires the correct maturation of Pit by Hsp90. Further research is necessary to understand how their chaperones and co-chaperones orchestrate OsCERK1 and Pit.

## Materials and Methods

### Plasmid constructs

The cDNAs of *Pit, OsCERK1*, and *OsRac1* were described previously (Chen et al. 2010a; Kawano et al. 2010a). They were transferred into various vectors, depending on the experiment. These included pBI221-Vn-Gateway, pBI221-Gateway-Vc (provided by Dr. Seiji Takayama, University of Tokyo), and 35S-Gateway-Venus/GFP. We generated three pZH2B vectors containing the *Ubiquitin* promoter (*UbqPro*)-*4×myc-OsRac1 wild type* (*WT*)-*NOS* terminator (*NOSTer*)-*UbqPro-OsCERK1-3×FLAG-NosTer* (Fig. 2A), *Ubq*Pro-*4×myc-OsRac1 WT*-*NOS*Ter-*Ubq*Pro-Pit WT-3*×FLAG*-*Nos*Ter (Fig. 2B), and *UbqPro-4×myc-OsRac1 WT-NOSTer-UbqPro-OsCERK1-3×FLAG-NosTer-Ubq*Pro-Pit*×FLAG-Nos*Ter (Fig. 1A) by multiple steps of PCR, subcloning, and enzymatic digestion. We also produced two estradiol-inducible vectors pER8-*Pit WT* and *D485V-3×FLAG* with *UbqPro-4×myc-OsRac1-NosTer* (Fig. 4) (pER8 was provided by Dr. Nam-Hai Chua, Rockefeller University). pGPVX:Hsp90 (10-186) was generated using pGPVX:Hsp90 (10-186) vector (provided by Dr. Ken-Ichiro Taoka, Yokohama City University) containing the backbone of pGreen (Hellens et al. 2000), 35S promotor and PVX region of piX.erG3 (Tamai and Meshi 2001), and Hsp90 (10-186) (Lu et al. 2003).

### Transgenic plants

Rice (*Oryza sativa* L. cv. Kinmaze) was used as the wild type and parental cultivar for the transgenic studies. Transgenic rice plants were generated using *Agrobacterium*-mediated transformation of rice calli (Hiei et al. 1994), and hygromycin-resistant plants were regenerated from transformed callus.

### Immunoprecipitation assay

For co-IP assay, 500 mgof rice cultured suspension cells frozen in liquid nitrogen or rice leaf blade samples were homogenized using a mortar with 1 ml of protein extraction buffer [50 mM Tris, pH 7.5, 2 mM EDTA, 150 mM NaCl, 5 mM MgCl2, 10% glycerol, 0.8% (w/v) Triton X-100, 1× protease inhibitor cocktail, and phosphatase inhibitor cocktail 1 and 2 (Sigma)]. After a 20-min incubation on ice, the homogenized samples were centrifuged at 20,000 × *g* for 20 min, and the resultant supernatants were collected. Using the BCA Protein Assay Reagent (Pierce), the protein concentration of the supernatants was measured and adjusted to 5 mg/ml protein with the protein extraction buffer. For co-IP of Myc-tagged and HA-tagged proteins, the μMACS c-myc Isolation Kit and μMACS HA Isolation Kit (Miltenyi Biotec) were used according to the instructions provided. For co-IP of FLAG-tag proteins, Immunoprecipitation Kit-Dynabeads protein G (Invitrogen) and Anti-FLAG M2 Monoclonal Antibody (Sigma-Aldrich) were used according to the instructions provided.

### BiFC assay

For use in BiFC experiments, *OsCERK1, Pit* and *OsRac1, Hop/Sti1a*, and *OsFLS2* were cloned into BiFC vectors, which were then purified using the Purelink Plasmid Midiprep Kit (Invitrogen) and introduced into rice protoplasts as described previously (Kawano et al. 2014a; Wong et al. 2018). The mCherry expression plasmid was introduced simultaneously as a marker for transformed cells. BiFC images were acquired using a TCS SP5 confocal microscope (Leica).

### VIAFM observation

*Pit* and *OsCERK1* were cloned into the *p35S-Gateway-mEGFP* and *-mCherry* vectors, respectively, for C-terminal fusion using LR reactions (Thermo Fisher Scientific). Rice protoplasts were transformed with these vectors as described previously (Wong et al. 2018). Ten to twelve hours after transformation, the cells were placed on a cover glass (25 × 60 mm, NO.1; Matsunami) and then covered with another cover glass (25 × 40 mm, NO.1; Matsunami) thinly coated with low gelling temperature agarose (Sigma, cat. no. A9414). VIAFM images were acquired using an Olympus TIRF system based on an Olympus IX81 equipped with an APON 60XO TIRF (N.A.: 1.49). mEGFP and mCherry were excited with 488- and 561-nm lasers, respectively.

### Subcellular localization in rice protoplasts

Venus was fused to either the C or the N terminus of Pit using the Gateway system (Invitrogen). The Pit-Venus, mCherry, and Cerulean-NLS constructs were controlled by the CaMV 35S promoter. Protoplast isolation from rice Oc suspension cultures and protoplast transformations were performed as described (Wong et al. 2018). Some of the transfected cells were treated with BFA (50 μg/ml: Sigma) and GDA (10 μM: Sigma). After incubation for 16 h at 30°C, the protoplasts were observed with a Leica TCS-SP5 microscope.

### Gel filtration

One hundred fifty milligrams of rice cell culture was ground in liquid nitrogen and extracted in 1 ml of protein extraction buffer for 20 min at 4°C. The extracts were centrifuged at 20,000 × *g* for 20 min at 4°C, and the supernatant was filtered through a 0.22-μm filter (Millipore). The filtrate was applied to a Superdex 200 column (GE Healthcare) attached to an AKTA Explorer system (GE Healthcare) using protein extraction buffer as the running buffer. LMW and HMW Gel Filtration calibration kits (GE Healthcare) were used to estimate the molecular weight of protein complexes. Fractions of 0.5 ml each were collected and 45-μl aliquots were concentrated by TCA/acetone precipitation. The precipitate was dissolved in 15 μl of SDS-PAGE sample buffer and treated for 20 min at 60°C. These samples were subjected to SDS-PAGE and immunoblot analysis.

### Pull-down assay using PAK CRIB

Purified GST-PAK CRIB was prepared according to a previous method (Kawano et al. 2010a). Rice cell cultures were ground in liquid nitrogen and extracted in 1 ml of protein extraction buffer for 20 min at 4°C. The extracts were centrifuged at 20,000 × *g* for 20 min at 4°C, and the supernatant was collected. Protein content was determined by the BCA assay reagent (Thermo Fisher Scientific), using bovine serum albumin (BSA) as a standard. Three milligrams of the total protein samples were applied to 20 μg of GST-PAK-CRIB glutathione Sepharose 4B for pulldown assays and rotated for 30 min at 4°C. The Sepharose was washed three times in protein extraction buffer. Proteins that remained bound to the Sepharose were eluted in 80 μl of SDS-PAGE sample buffer and treated for 20 min at 60°C. These samples were subjected to SDS-PAGE, immunoblot analysis, and Coomassie staining.

### Preparation of membrane fractions

Rice cell cultures were harvested 3 days after subculture and homogenized in homogenizing medium [50 mM MOPS/KOH, pH 7.6, 5 mM EGTA, 5 mM EDTA, 0.5 M D-sorbitol, 1.5% (w/v) polyvinylpyrrolidone, 2 mM PMSF, 2.5 mM DTT]. The homogenate was filtered through Miracloth (Calbiochem), and the filtrate was centrifuged at 3,000 × *g* for 10 min at 4°C. The supernatant was collected and centrifuged at 170,000 × *g* for 35 min at 4°C to yield soluble (supernatant) and microsomal (pellet) protein fractions. A polyethylene glycol–dextran (6.4%, w/w) aqueous two-phase partitioning system (Fujiwara et al. 2009) was used to separate the plasma membrane (PM) and endomembranes (EMs). The microsomal pellets were resuspended in MS-suspension medium [10 mM potassium phosphate, pH 7.8, 300 mM sucrose] and subjected to two-phase partitioning. Both the upper phase (enriched for the PM) and the lower phase (enriched for the EMs) were partitioned three times with lower phase buffer and upper phase buffer, respectively. The PM and EM fractions were harvested by centrifugation at 170,000 × *g* for 35 min at 4°C, and resuspended in PM-suspension medium (10 mM MOPS/KOH, pH 7.3, 1 mM EGTA, 300 mM sucrose, 2 mM DTT). The protein content of the fractions was determined by the BCA assay reagent (Thermo Fisher Scientific), using BSA as a standard. These samples were subjected to SDS-PAGE and immunoblot analysis.

### Immunoblotting

Sample proteins were separated by SDS-PAGE and electrotransferred onto an Immobilon-P membrane (Millipore) for immunoblot detection. The membrane was blocked for 1 h in Blocking One (Nacalai Tesque) for 30 min and incubated for 30 min with anti-Myc (Nacalai Tesque) or anti-RACK1A (Nakashima et al. 2008), anti-FLAG (Sigma), anti-Hop/Sti1a (Chen et al. 2010a), anti-OsCEBiP (Kaku et al. 2006), anti-Hsp90 (Enzo Life Sciences), anti-SGT1 (Azevedo et al. 2002), anti-RAR1 (Thao et al. 2007), anti-OsMPK6 (Lieberherr et al. 2005), anti-tubulin (Calbiochem), anti-Bip (Cosmo Bio), and anti-OsPIP1s (Cosmo Bio) antibodies. After washing twice with TBST (0.05 M Tris, pH 7.6, 0.9% NaCl, 0.1% Triton X-100), the membranes were incubated for 2 h in Can Get Signal Solution 2 (Toyobo) with anti-rabbit or mouse IgG conjugated to horseradish peroxidase (GE Healthcare). After washing twice with TBST, chemical enhancement was performed using ECL PLUS Western blot detection reagents (GE Healthcare). The enhanced signals were detected by the LAS-4000 system (Fujifilm).

### Agroinfiltration into *N. benthamiana* leaves

In some experiments, we generated *pGPVX:Hsp90* (10-186) vector and virus-induced gene silencing (VIGS) was done as described by Lu *et al*. (Lu et al. 2003). Agroinfiltration of *N. benthamiana* was performed as described previously (Kawano et al. 2010a). *Agrobacterium tumefaciens* strain GV3010, harboring the helper plasmid pSoup and binary plasmids carrying the cDNAs of Pit WT and mutants, was used to infiltrate leaves of 5-week-old *N. benthamiana* plants. We used the p19 silencing suppressor to enhance gene expression. Each *Agrobacterium* culture was resuspended in a buffer containing 10 mM MgCl_2_, 10 mM MES, pH 5.6, and 150 μM acetosyringone, and incubated at 23°C for 2–3 h before infiltration. In some experiments, we added GDA at a final concentration of 10 μM to an *Agrobacterium* culture carrying pGWB2-*Pit D485V*. The plants were kept in a growth chamber at 23°C after agroinfiltration. To visualize hydrogen peroxide, a major endogenous ROS, *in situ*, the agroinfiltrated leaves were detached and incubated in 1 μg/ml DAB solution for 2–8 h, after which they were decolorized in boiling ethanol. Photographs were taken at 7 days post-inoculation (dpi) for cell death at 3 dpi for ROS production.

**Table1.**
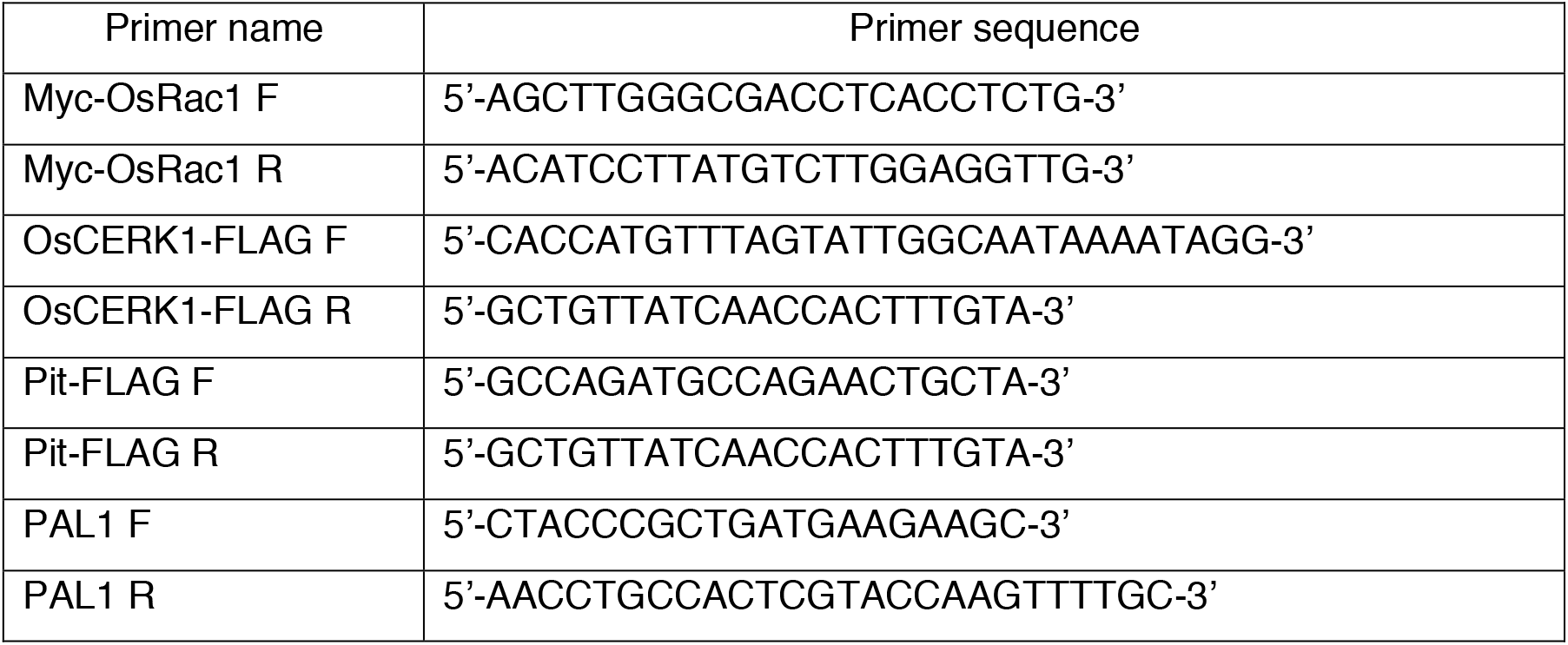
List of primers.

## Acknowledgments

We thank the members of the Laboratory of Plant Molecular Genetics at NAIST, the Laboratory of Signal Transduction and Immunity at PSC, and the Plant Immune Design Group at Okayama University for invaluable support and discussions.

---

**Supplementary Figure 1.**
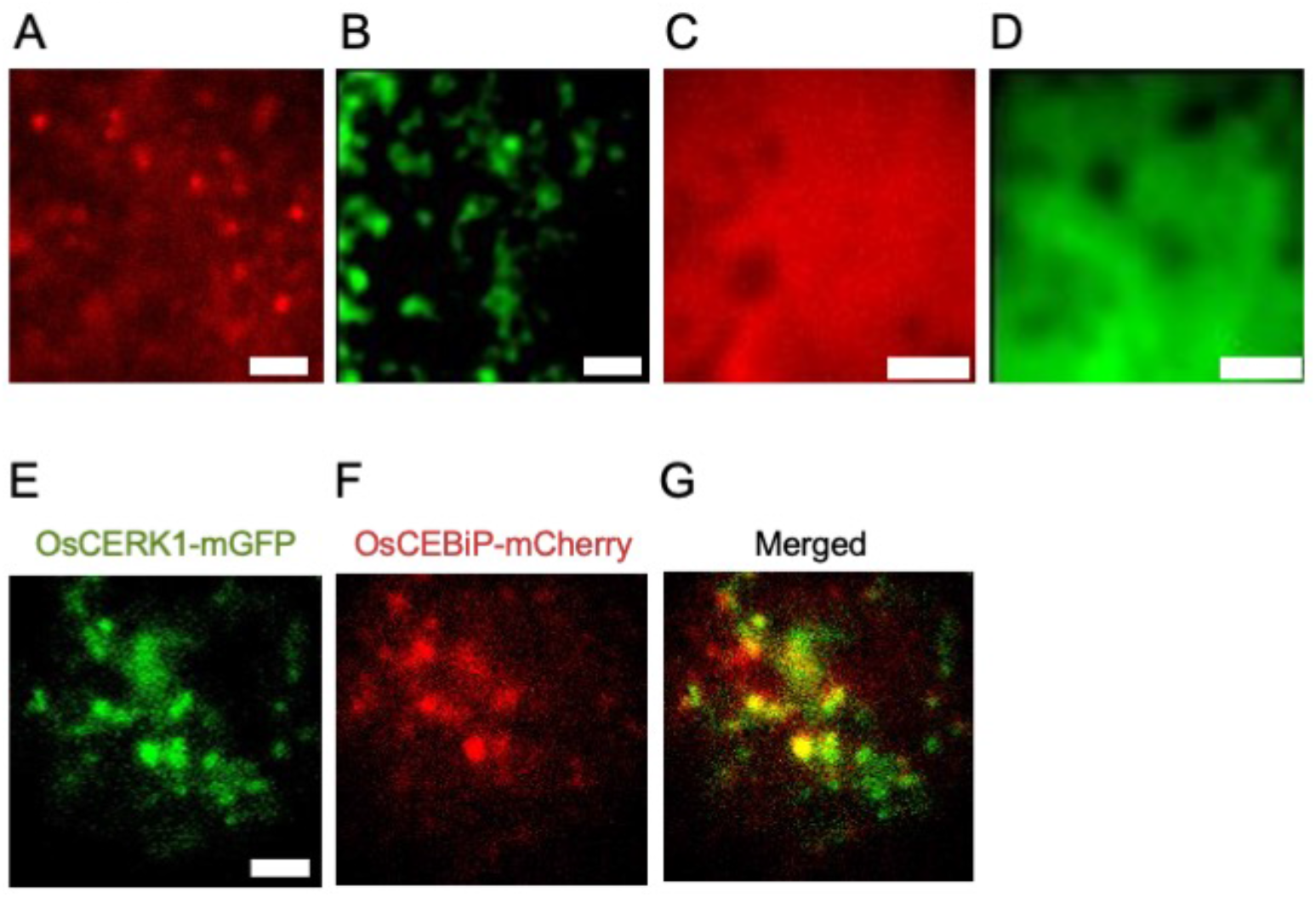
VIAFM images of the immune receptors. Representative image of OsCERKI-mCherry (A), Pit-mEGFP (B), mCherry (C), and mEGFP (D). E-F, co-expression of 0sCERK1-mGFP and OsCEBiP-mCherry. Scale bar is 5 μm.

## References

Akamatsu, A., Uno, K., Kato, M., Wong, H.L., Shimamoto, K. and Kawano, Y. (2015) New insights into the dimerization of small GTPase Rac/ROP guanine nucleotide exchange factors in rice. Plant Signal Behav 10: e1044702.

Akamatsu, A., Wong, H., Fujiwara, M., Okuda, J., Nishide, K., Uno, K., et al. (2013) An OsCEBiP/OsCERK1-OsRacGEF1-OsRac1 module is an essential component of chitin-induced rice immunity. Cell Host Microbe 13: 465–476.

Azevedo, C., Sadanandom, A., Kitagawa, K., Freialdenhoven, A., Shirasu, K. and Schulze-Lefert, P. (2002) The RAR1 interactor SGT1, an essential component of R gene-triggered disease resistance. Science 295: 2073–2076.

Berken, A. (2006) ROPs in the spotlight of plant signal transduction. Cell Mol Life Sci 63: 2446–2459.

Bieri, S., Mauch, S., Shen, Q.H., Peart, J., Devoto, A., Casais, C., et al. (2004) RAR1 positively controls steady state levels of barley MLA resistance proteins and enables sufficient MLA6 accumulation for effective resistance. Plant Cell 16: 3480–3495.

Chen, L., Hamada, S., Fujiwara, M., Zhu, T., Thao, N.P., Wong, H.L., et al. (2010a) The Hop/Sti1-Hsp90 chaperone complex facilitates the maturation and transport of a PAMP receptor in rice innate immunity. Cell Host Microbe 7: 185–196.

Chen, L., Shiotani, K., Togashi, T., Miki, D., Aoyama, M., Wong, H.L., et al. (2010b) Analysis of the Rac/Rop small GTPase family in rice: expression, subcellular localization and role in disease resistance. Plant Cell Physiol 51: 585–595.

Couto, D. and Zipfel, C. (2016) Regulation of pattern recognition receptor signalling in plants. Nat Rev Immunol 16: 537–552.

Cui, H., Tsuda, K. and Parker, J.E. (2015) Effector-triggered immunity: from pathogen perception to robust defense. Annu Rev Plant Biol 66: 487–511.

Dangl, J.L., Horvath, D.M. and Staskawicz, B.J. (2013) Pivoting the plant immune system from dissection to deployment. Science 341: 746–751.

Dodds, P.N. and Rathjen, J.P. (2010) Plant immunity: towards an integrated view of plant-pathogen interactions. Nat Rev Genet 11: 539–548.

Fujimoto, M., Arimura, S., Ueda, T., Takanashi, H., Hayashi, Y., Nakano, A., et al. (2010) Arabidopsis dynamin-related proteins DRP2B and DRP1A participate together in clathrin-coated vesicle formation during endocytosis. Proc Natl Acad Sci U S A 107: 6094–6099.

Fujiwara, M., Hamada, S., Hiratsuka, M., Fukao, Y., Kawasaki, T. and Shimamoto, K. (2009) Proteome analysis of detergent-resistant membranes (DRMs) associated with OsRac1-mediated innate immunity in rice. Plant Cell Physiol 50: 1191–1200.

Hellens, R.P., Edwards, E.A., Leyland, N.R., Bean, S. and Mullineaux, P.M. (2000) pGreen: a versatile and flexible binary Ti vector for Agrobacterium-mediated plant transformation. Plant Mol Biol 42: 819–832.

Hemsley, P.A. (2015) The importance of lipid modified proteins in plants. New Phytol 205: 476–489.

Hiei, Y., Ohta, S., Komari, T. and Kumashiro, T. (1994) Efficient transformation of rice (Oryza sativa L.) mediated by Agrobacterium and sequence analysis of the boundaries of the T-DNA. Plant J 6: 271–282.

Hubert, D.A., Tornero, P., Belkhadir, Y., Krishna, P., Takahashi, A., Shirasu, K., et al. (2003) Cytosolic HSP90 associates with and modulates the Arabidopsis RPM1 disease resistance protein. EMBO J 22: 5679–5689.

Ishiguro, S., Watanabe, Y., Ito, N., Nonaka, H., Takeda, N., Sakai, T., et al. (2002) SHEPHERD is the Arabidopsis GRP94 responsible for the formation of functional CLAVATA proteins. EMBO J 21: 898–908.

Kadota, Y. and Shirasu, K. (2012) The HSP90 complex of plants. Biochim Biophys Acta 1823: 689–697.

Kadota, Y., Shirasu, K. and Guerois, R. (2010) NLR sensors meet at the SGT1-HSP90 crossroad. Trends Biochem Sci 35: 199–207.

Kaku, H., Nishizawa, Y., Ishii-Minami, N., Akimoto-Tomiyama, C., Dohmae, N., Takio, K., et al. (2006) Plant cells recognize chitin fragments for defense signaling through a plasma membrane receptor. Proc Natl Acad Sci U S A 103: 11086–11091.

Kawano, Y., Akamatsu, A., Hayashi, K., Housen, Y., Okuda, J., Yao, A., et al. (2010a) Activation of a Rac GTPase by the NLR family disease resistance protein Pit plays a critical role in rice innate immunity. Cell Host Microbe 7: 362–375.

Kawano, Y., Chen, L. and Shimamoto, K. (2010b) The function of Rac small GTPase and associated proteins in rice innate immunity. Rice 3: 112–121.

Kawano, Y., Fujiwara, T., Yao, A., Housen, Y., Hayashi, K. and Shimamoto, K. (2014a) Palmitoylation-dependent membrane localization of the rice resistance protein pit is critical for the activation of the small GTPase OsRac1. J Biol Chem 289: 19079–19088.

Kawano, Y., Kaneko-Kawano, T. and Shimamoto, K. (2014b) Rho family GTPase-dependent immunity in plants and animals. Front in Plant Sci 5: 522.

Kawano, Y. and Shimamoto, K. (2013) Early signaling network in rice PRR- and R-mediated immunity. Curr Opin Plant Biol 16: 496–504.

Kawasaki, T., Henmi, K., Ono, E., Hatakeyama, S., Iwano, M., Satoh, H., et al. (1999) The small GTP-binding protein Rac is a regulator of cell death in plants. Proc Natl Acad Sci U S A 96: 10922–10926.

Kawasaki, T., Koita, H., Nakatsubo, T., Hasegawa, K., Wakabayashi, K., Takahashi, H., et al. (2006) Cinnamoyl-CoA reductase, a key enzyme in lignin biosynthesis, is an effector of small GTPase Rac in defense signaling in rice. Proc Natl Acad Sci U S A 103: 230–235.

Kim, S.H., Oikawa, T., Kyozuka, J., Wong, H.L., Umemura, K., Kishi-Kaboshi, M., et al. (2012) The bHLH Rac immunity1 (RAI1) is activated by OsRac1 via OsMAPK3 and OsMAPK6 in rice immunity. Plant Cell Physiol 53: 740–754.

Konopka, C.A. and Bednarek, S.Y. (2008) Variable-angle epifluorescence microscopy: a new way to look at protein dynamics in the plant cell cortex. Plant J 53: 186–196.

Kosami, K., Ohki, I., Nagano, M., Furuita, K., Sugiki, T., Kawano, Y., et al. (2014) The crystal structure of the plant small GTPase OsRac1 reveals its mode of binding to NADPH oxidase. J Biol Chem 289: 28569–28578.

Leister, R.T., Dahlbeck, D., Day, B., Li, Y., Chesnokova, O. and Staskawicz, B.J. (2005) Molecular genetic evidence for the role of SGT1 in the intramolecular complementation of Bs2 protein activity in Nicotiana benthamiana. Plant Cell 17: 1268–1278.

Lieberherr, D., Thao, N.P., Nakashima, A., Umemura, K., Kawasaki, T. and Shimamoto, K. (2005) A sphingolipid elicitor-inducible mitogen-activated protein kinase is regulated by the small GTPase OsRac1 and heterotrimeric G-protein in rice. Plant Physiol 138: 1644–1652.

Liu, Y., Burch-Smith, T., Schiff, M., Feng, S. and Dinesh-Kumar, S.P. (2004) Molecular chaperone Hsp90 associates with resistance protein N and its signaling proteins SGT1 and Rar1 to modulate an innate immune response in plants. J Biol Chem 279: 2101–2108.

Lu, R., Malcuit, I., Moffett, P., Ruiz, M.T., Peart, J., Wu, A.J., et al. (2003) High throughput virus-induced gene silencing implicates heat shock protein 90 in plant disease resistance. EMBO J 22: 5690–5699.

Monaghan, J. and Zipfel, C. (2012) Plant pattern recognition receptor complexes at the plasma membrane. Curr Opin Plant Biol 15: 349–357.

N’gou, B.P.M., Ahn, H.-K., Ding, P. and Jones, J. (2020) Mutual Potentiation of Plant Immunity by Cell-surface and Intracellular Receptors. bioRxiv https://doi.org/10.1101/2020.04.10.034173.

Nagano, M., Ishikawa, T., Fujiwara, M., Fukao, Y., Kawano, Y., Kawai-Yamada, M., et al. (2016) Plasma Membrane Microdomains Are Essential for Rac1-RbohB/H-Mediated Immunity in Rice. Plant Cell 28: 1966–1983.

Nakashima, A., Chen, L., Thao, N.P., Fujiwara, M., Wong, H.L., Kuwano, M., et al. (2008) RACK1 functions in rice innate immunity by interacting with the Rac1 immune complex. Plant Cell 20: 2265–2279.

Nibau, C., Wu, H.M. and Cheung, A.Y. (2006) RAC/ROP GTPases: ‘hubs’ for signal integration and diversification in plants. Trends Plant Sci 11: 309–315.

Ono, E., Wong, H.L., Kawasaki, T., Hasegawa, M., Kodama, O. and Shimamoto, K. (2001) Essential role of the small GTPase Rac in disease resistance of rice. Proc Natl Acad Sci U S A 98: 759–764.

Pearl, L.H. and Prodromou, C. (2006) Structure and mechanism of the Hsp90 molecular chaperone machinery. Annu Rev Biochem 75: 271–294.

Peng, Y., van Wersch, R. and Zhang, Y. (2018) Convergent and Divergent Signaling in PAMP-Triggered Immunity and Effector-Triggered Immunity. Mol Plant Microbe Interact 31: 403–409.

Qi, Y., Tsuda, K., Glazebrook, J. and Katagiri, F. (2011) Physical association of pattern-triggered immunity (PTI) and effector-triggered immunity (ETI) immune receptors in Arabidopsis. Mol Plant Pathol 12: 702–708.

Sander, E.E., van Delft, S., ten Klooster, J.P., Reid, T., van der Kammen, R.A., Michiels, F., et al. (1998) Matrix-dependent Tiam1/Rac signaling in epithelial cells promotes either cell-cell adhesion or cell migration and is regulated by phosphatidylinositol 3-kinase. J Cell Biol 143: 1385–1398.

Shirasu, K., Lahaye, T., Tan, M.W., Zhou, F., Azevedo, C. and Schulze-Lefert, P. (1999) A novel class of eukaryotic zinc-binding proteins is required for disease resistance signaling in barley and development in C. elegans. Cell 99: 355–366.

Takahashi, A., Casais, C., Ichimura, K. and Shirasu, K. (2003) HSP90 interacts with RAR1 and SGT1 and is essential for RPS2-mediated disease resistance in Arabidopsis. Proc Natl Acad Sci U S A 100: 11777–11782.

Takeuchi, M., Ueda, T., Sato, K., Abe, H., Nagata, T. and Nakano, A. (2000) A dominant negative mutant of sar1 GTPase inhibits protein transport from the endoplasmic reticulum to the Golgi apparatus in tobacco and Arabidopsis cultured cells. Plant J 23: 517–525.

Tamai, A. and Meshi, T. (2001) Cell-to-cell movement of Potato virus X: the role of p12 and p8 encoded by the second and third open reading frames of the triple gene block. Mol Plant Microbe Interact 14: 1158–1167.

Tao, Y., Xie, Z., Chen, W., Glazebrook, J., Chang, H.S., Han, B., et al. (2003) Quantitative nature of Arabidopsis responses during compatible and incompatible interactions with the bacterial pathogen Pseudomonas syringae. Plant Cell 15: 317–330.

Thao, N.P., Chen, L., Nakashima, A., Hara, S., Umemura, K., Takahashi, A., et al. (2007) RAR1 and HSP90 form a complex with Rac/Rop GTPase and function in innate-immune responses in rice. Plant Cell 19: 4035–4045.

Thomma, B.P., Nurnberger, T. and Joosten, M.H. (2011) Of PAMPs and effectors: the blurred PTI-ETI dichotomy. Plant Cell 23: 4–15.

Tsuda, K. and Katagiri, F. (2010) Comparing signaling mechanisms engaged in pattern-triggered and effector-triggered immunity. Curr Opin Plant Biol 13: 459–465.

Tsuda, K., Mine, A., Bethke, G., Igarashi, D., Botanga, C.J., Tsuda, Y., et al. (2013) Dual regulation of gene expression mediated by extended MAPK activation and salicylic acid contributes to robust innate immunity in Arabidopsis thaliana. PLoS genetics 9: e1004015.

Wang, Q., Li, Y., Ishikawa, K., Kosami, K.I., Uno, K., Nagawa, S., et al. (2018) Resistance protein Pit interacts with the GEF OsSPK1 to activate OsRac1 and trigger rice immunity. Proc Natl Acad Sci U S A 115: E11551–E11560.

Wong, H.L., Akamatsu, A., Wang, Q., Higuchi, M., Matsuda, T., Okuda, J., et al. (2018) In vivo monitoring of plant small GTPase activation using a Forster resonance energy transfer biosensor. Plant Methods 14: 56.

Wong, H.L., Pinontoan, R., Hayashi, K., Tabata, R., Yaeno, T., Hasegawa, K., et al. (2007) Regulation of rice NADPH oxidase by binding of Rac GTPase to its N-terminal extension. Plant Cell 19: 4022–4034.

Yalovsky, S., Bloch, D., Sorek, N. and Kost, B. (2008) Regulation of membrane trafficking, cytoskeleton dynamics, and cell polarity by ROP/RAC GTPases. Plant Physiol 147: 1527–1543.

Yuan, M., Jian, Z., Bi, G., Nomura, K., Liu, M., He, S.Y., et al. (2020) Pattern-recognition receptors are required for NLR-mediated plant immunity. bioRxiv https://doi.org/10.1101/2020.04.10.031294.

